# Enrichment of neurodegenerative microglia signature in brain-derived extracellular vesicles isolated from Alzheimer’s disease mouse model

**DOI:** 10.1101/2020.11.05.369728

**Authors:** Satoshi Muraoka, Mark P. Jedrychowski, Naotoshi Iwahara, Mohammad Abdullah, Kristen D. Onos, Kelly J. Keezer, Jianqiao Hu, Seiko Ikezu, Gareth R. Howell, Steven P. Gygi, Tsuneya Ikezu

## Abstract

Extracellular vesicles (EVs) are secreted by any neuronal cells in the central nervous system (CNS) for molecular clearance, cellular communications and disease spread in multiple neurodegenerative diseases, including Alzheimer’s disease (AD), although their exact molecular mechanism is poorly understood. We hypothesize that high-resolution proteomic profiling of EVs separated from animal models of AD would determine the composition of EV contents and their cellular origin. Here, we examined recently developed transgenic mice (CAST.*APP/PS1*), which express familial AD-linked mutations of amyloid precursor protein (*APP*) and presenilin-1 (*PS1*) in the CAST/EiJ mouse strain and develop hippocampal neurodegeneration. Quantitative proteomics analysis of EVs separated from CAST.*APP/PS1* and age-matched control mice by tandem mass tag-mass spectrometry identified a total of 3,444 unique proteins, which are enriched in neuron, astrocyte, oligodendrocyte and microglia-specific molecules. CAST.*APP/PS1*-derived EVs show significant enrichment of Psen1, APP, Itgax, and reduction of Wdr61, Pmpca, Aldh1a2, Calu, Anp32b, Actn4 and Ndufv2 compared to WT-derived EVs, suggesting the involvement of Aβ-processing complex and disease-associated / neurodegenerative microglia (DAM/MGnD) in EV secretion. In addition, Itgax and Apoe, the DAM/MGnD markers, in EV show a positive correlation with *Itgax* and *Apoe* mRNA expression from brain tissue in CAST.*APP/PS1* mice. These datasets indicate the significant contribution of Aβ plaque and neurodegeneration-induced DAM/MGnD microglia for EV secretion in CAST.*APP/PS1* mice and shed light on understanding the AD pathogenesis.

## Introduction

Alzheimer’s disease (AD) is a progressive neurodegenerative disorder and the most common forms of adult dementia affecting 50 million people worldwide (1). The neuropathology of AD is characterized by extracellular deposition of amyloid-β (Aβ) plaques, which are processed by amyloid precursor protein (APP) and presenilin-1 (PS1)-dependent gamma secretase complex, and intraneuronal accumulation of neurofibrillary tangles (NFTs), which are consisted with hyperphosphorylated microtubule-associated protein tau (2–4). There are two form of AD, early-onset / familial AD (FAD) and sporadic / late-onset AD (LOAD) (5, 6). FAD is mostly caused by mutations in *APP* and *PSEN1* and *PSEN2* (7). The FAD mouse models expressing FAD-linked mutation of *APP, PSEN1* or both, have been extensively used to understand the pathophysiology of Aβ deposition although most of them do not develop neurodegeneration (8–10). Onos *et al.* have recently reported a comprehensive assessment of the transgene expression of FAD-linked mutation of *APP* and *PSEN1* in different genetic backgrounds including B6, WSB/EiJ, PWK/PhJ, and CAST/EiJ to establish more clinically-relevant AD mouse models (11). The study showed that CAST.*APP/PS1* line develops reduction in the number of hippocampal pyramidal neurons and robust neuroinflammatory response than previous models (11), which would be more suitable for the assessment of Aβ deposition-induced inflammatory reaction and neuronal cell loss.

Extracellular vesicles (EVs), including exosomes (50-150nm), ectosomes/microvesicles (150-1000nm), and apoptotic bodies (1000-5000nm) are released from almost any neuronal cells (12–14). These EVs contain proteins, mRNA, non-coding RNAs (such as microRNA) and lipids, can transfer these molecules from cells to cells, and can be transported to biofluids, such as and cerebrospinal fluid and blood. In the central nervous system (CNS), brain-derived EVs contain multiple AD-associated proteins such as Aβ α-synuclein, APP, cyclin-dependent kinase 5, PSEN1, and tau, and play important roles in Aβ deposition and tauopathy (15–20). Moreover, it has been reported that inhibition of EV synthesis reduced Aβ plaque deposition in the mouse model of AD, and stimulation of EV secretion increased intracellular transfer of prion protein in AD mouse models (16, 17). EVs are involved in the extracellular enzymatic degradation of Aβ and promote both Aβ aggregation and clearance by microglia (18, 19), although their exact molecular mechanism is poorly understood. We hypothesize that high-resolution proteomic profiling of EVs separated from animal models of AD would determine the composition of EV contents and their cellular origin. Here we provide the quantitative proteomics profiling of EVs separated from CAST.*APP/PS1* transgenic mouse brain tissue and show brain-derived EV molecules altered during early-onset AD.

## Materials and methods

### CAST.-*APP/PS1* transgenic mouse model

The CAST.-*APP/PS1* transgenic mouse line, which expresses human *APP*^*swe*^ and *PS1*^*de9*^, was created in the Howell lab colony at The Jackson Laboratory by backcrossing for at least seven generations the *APP/PS1* transgenes from C57BL/6J (B6) to CAST (11). Brain samples (forebrain and hindbrain) were extracted from 6 female CAST.*APP/PS1* and 6 female CAST (WT) littermate control mice at 8 months of age. Mice were anesthetized with ketamine/xylazine prior to tissue harvest.

### Brain tissue homogenates

Frozen whole brain tissue was chopped on ice using a razor blade (# 12-640 Fischer Scientific) to generate approximately 0.5 mm-wide pieces, and homogenized by a sonicator. The homogenized tissue was lyzed using Guanidine Hydrochloride (# 50950-250G Sigma).

### Separation of EVs from mouse brain tissue

Brain tissue (0.4 g per sample) was processed for EV extraction based on our reported method with modifications. Briefly, frozen whole brain tissue was chopped on ice using a razor blade (# 12-640 Fischer Scientific) to generate approximately 0.5 mm-wide pieces. The sections were transferred to 3mL of Hibernate E solution (# A1247601 Gibco) containing 20 U of papain (# LK003178 Worthington-biochemical corporation) in Earle’s Balanced Salt Solution (EBSS) (# 14155063 Gibco) and then incubated at 37°C for 15 min by stirring once every 5 min. After the incubation, the samples were placed on ice, and added with 6 mL of ice-cold Hibernate E solution supplemented with Halt™ Protease and Phosphatase Inhibitor Cocktails (# PI78443 Fisher scientific). The samples were gently homogenized (20 strokes) with a glass-Teflon homogenizer (# 89026-384 VWR), and filtered with 40-μm mesh filter (# 22-363-547 Fisher scientific), followed by centrifugation at 300 × *g* for 10 min at 4°C (# 5720R Eppendorf). The supernatant was transferred to a new 15-mL polypropylene tube and centrifuged at 2,000 × *g* for 10 min at 4°C (# 5720R Eppendorf). The supernatant was transferred to a 30-mL conical tube and centrifuged at 10,000 × *g* for 10 min at 4°C (#5424R Eppendorf). The supernatant filtered through a 0.22-μm polyethersulfone membrane filter (# SLGP033RS EMD Millipore) into new a polyallomer ultracentrifuge tube with 13.2-mL capacity (# 331372 Beckman Coulter), diluted with double-filtered phosphate-buffered saline (dfPBS) with 0.22-μm polyethersulfone membrane filter to 12 mL, and centrifuged at 140,000 × *g* for 70 min at 4°C (# Optima-XE SW41 Beckman Coulter). The pellet was resuspended in 2 mL of 0.475M of sucrose solution (# S5-3 Fisher science) in dfPBS. The sucrose step gradient was created in dfPBS with six 2-mL steps starting from 2.0M to 1.5M, 1.0M, 0.825M, 0.65M, and 0.475M (containing the resuspended pellet) in a polyallomer ultracentrifuge tube. The gradient was centrifuged at 200,000 × *g* for 20 h at 4°C (35,000 rpm with # Optima-XE SW41 Beckman Coulter). The gradient was collected in 2-mL fractions, except for the first and last fractions, which were 1 mL each. The interphases between the second (0.65M) and third (0.825M) steps correspond to fraction “V” and the third and fourth steps corresponded to fraction “VI” have a buoyant density of 1.10 - 1.12 and 1.12 - 1.15 g/cm^3^, respectively, and enriched in EVs. The V and VI fractions were diluted to a total volume of 12 mL with dfPBS and centrifuged at 140,000 × *g* for 70 min at 4°C (# Optima-XE SW41 Beckman Coulter), and each pellet were resuspend with 30 μl of dfPBS. The fraction V and VI fractions were mixed as an EV-enriched sample.

### Protein concentrations

The bicinchoninic acid (BCA) assay was used to determine protein concentration for each sample using BCA protein assay kit (# 23225 Pierce) as previously described (21). EVs were diluted 1:10 before loading into the assay, and a 1:8 ratio of sample to reaction components was used. All assays were allowed to incubate at 37°C for 30 min before protein concentration was read in at 562 nm (SynergyMix, Biotek).

### Nanoparticle Tracking Analysis (NTA)

All samples were diluted in dfPBS at least 1:8000 to get particles within the target reading range for the Nanosight 300 machine (Malvern Panalytical Inc), which is 10-100 particles per frame. Using a manual injection system, four 60-s videos were taken for each sample at 21°C. Analysis of particle counts was carried out in the Nanosight NTA 3.2 software (Malvern Panalytical Inc) with a detection threshold of 5.

### Transmission electron microscopy (TEM)

The EV separated from *APP/PS1* and control mouse brain tissue were analyzed by TEM. The EV sample (5μl) was adsorbed for 1 min to a carbon-coated mesh grid (# CF400-CU, Electron Microscopy Sciences) that had been made hydrophilic by a 20-s exposure to a glow discharge (25 mA). Excess liquid was removed with a filter paper (# 1 Whatman). The grid was then floated briefly on a drop of water (to wash away phosphate or salt), blotted on a filer paper, and then stained with 0.75% uranyl formate (# 22451 Electron Microscopy Sciences) for 30 s. After removing the excess uranyl formate, the grids were examined, and random fields were photographed using a JEOL 1200EX TEM with an AMT 2k CCD camera at the Electron Microscopy Facility, Harvard Medical School, Boston, MA.

### Western blotting

EV samples and brain tissue homogenate samples were run in a 4% to 20% gradient gel (# 4561093 Bio-Rad) and electro-transferred to Immobilon-P membrane, PVDF 0.45-μm (# 10344661 Fisher scientific). The membrane was blocked in freshly prepared 5% BSA diluted in TBS before being immunoblotted with specific primary antibodies (CD81; #EXOAB-CD81A-1 System Biosciences, GM130; #610822 Becton Dickinson, Cytochrome C; #11940T Cell Signaling Technology, ANXA5, ItgaX; #14-011485 eBioscience) or HRP-labeled primary antibodies (TSG101; # SC-7964 Santa Cruz Biotechnology). The membrane was incubated with HRP-labeled secondary antibodies (Santa Cruz Biotechnology) and scanned using the C300 digital chemiluminescent imager (Azure Biosystems).

### High-Resolution Liquid Chromatography-Tandem Mass-tag Mass spectrometry SDS-PAGE and In-gel digestion

Ice-cold 100% (w / v) trichloroacetic acid (TCA) (# T6399 Sigma-Aldrich) was added to the separated EV fraction to a final concentration of 20% of TCA, then the mixed sample was incubated for 30 min on ice and was centrifuged at 15,000 g for 20 min at 4°C. The pellet was then washed twice with ice-cold acetone (# 179124 Sigma-Aldrich). After drying, the pellet was resuspended in Laemmli sample buffer (# 1610747 Bio-Rad) with 5 mM dithiothreitol (# 43815 Sigma-Aldrich), reduced for 20 min at 65°C, and alkylated with 15 mM iodoacetamide (# I1149 Sigma-Aldrich) for 20 min at room temperature in the dark. Subsequently, the samples were run in a 4% to 20% gradient gel (# 4561096 Bio-Rad) until the dye front was 10 mm from the top of the gel. The gels were washed twice with distilled water, fixed with 100% Methanol, and stained with GelCode Blue Stain Reagent (# 24590 Thermo Fisher Scientific) for 16 hrs. Each lane was then individually removed from the gel. Gel pieces were then transferred to 1.5 mL tubes and destained twice using 50% acetonitrile (J. T. Baker, USA) in 25 mM HEPES (pH 8.8) at 22°C, for 15 min with shaking, and dehydrated with 100% acetonitrile for additional 10 min with shaking, for a total of three times. The destained gel piece was dried up using SpeedVac Concentrators (Thermo Fisher Scientific). The gel pieces were digested with proteomic grade trypsin (# 03708985 Roche, USA) in 25 mM HEPES overnight at 37°C. The digested peptide was extracted with 70% acetonitrile /1% formic acid, and were removed the gel by Ultrafree-MC Centrifugal Filter (# UFC30L Millipore USA). The digested peptides were reconstituted in 25 μL of 200 mM EPPS (pH 8.0) and vortexed for 5 min.

### Peptide labeling with TMT 16-plex isobaric labeling Kit

Tandem mass tag (TMT) labeling was performed according to manufacturer’s instructions (# A44520 Thermo Fisher Scientific). In brief, 4 μL of TMT Label reagent (20 ng/μL) was added to the digested peptides in 30 μL of 200 mM HEPPS (4-(2-Hydroxyethyl)-1-piperazinepropanesulfonic acid), pH8.0. After incubation at room temperature for 1 h, the reaction was quenched with 2 μL of 5% hydroxylamine in water for 15min. The TMT-labeled peptide samples were pooled at a 1:1 ratio across ten samples. The combined sample was added to 100 μL of 20% formic acid, 2 mL of 1% formic acid, desalted via StageTip, dried by vacuum centrifugation, and resuspended in 20 μL of 5% acetonitrile and 5% formic acid for nano liquid chromatography and tandem mass-spectrometry (Nano LC-MS/MS/MS).

### Nano-Liquid Chromatography and Tandem Mass-tag Spectrometry (LC-MS/MS/MS)

Nano LC–MS/MS/MS analysis was conducted using an LTQ-Orbitrap Fusion Lumos mass spectrometer (Thermo Fisher Scientific, USA) equipped with a Proxeon EASY-nano LC 1200 liquid chromatography pump (Thermo Fisher Scientific, San Jose, CA). Peptides were separated on a 100 μm inner diameter microcapillary column packed with 35-cm long Accucore150 resin (2.6 μm, 150 Å, Thermo Fisher Scientific). We loaded 4 μL onto the column and separation was achieved using a 180 min gradient of 8 to 23% acetonitrile in 0.125% formic acid at a flow rate of ~550 nL/min. The analysis used an MS^3^ based TMT method, which has been shown to reduce ion interference. The scan sequence began with an MS1 spectrum (Orbitrap; resolution 120,000; mass range 400-1400m/z; automatic gain control (AGC) target 5 × 10^5^; maximum injection time 100ms). Precursors for MS^2^/MS^3^ analysis were selected using a Top10 method. MS2 analysis consisted of collision-induced dissociation (quadrupole ion trap; AGC 2 × 10^4^; normalized collision energy (NCE) 35; maximum injection time 150ms). Following acquisition of each MS^2^ spectrum, we collected an MS^3^ spectrum using our recently described method in which multiple MS^2^ fragment ions were captured in the MS^3^ precursor population using isolation waveforms with multiple frequency notches (22). MS^3^ precursors were fragmented by high-energy collision-induced dissociation (HCD) and analyzed using the Orbitrap (NCE 65; AGC 1 × 10^5^; maximum injection time 150ms, resolution was 50,000 at 200Th).

### Mass-spectrometry data analysis

A compendium of in-house developed software was used to convert mass spectrometric data (Raw file) to the mzXML format, as well as to correct monoisotopic m/z measurements (23). Database searching included all entries from the *Mus musculus* with human APP and PS1 UniProt database (ver. October 2018). This database was concatenated with one composed of all protein sequences in the reversed order. Searches were performed using a 50ppm precursor ion tolerance for total protein level profiling (22). The product ion tolerance was set to 0.9 Da, which was chosen to maximize sensitivity in conjunction with SEQUEST searches and linear discriminant analysis. TMT tags on lysine residues and peptide N termini (+229.163 Da) and carbamidomethylation of cysteine residues (+57.021 Da) were set as static modifications, while oxidation of methionine residues (+15.995 Da) was set as a variable modification. Peptide-spectrum matches (PSMs) were adjusted to a 1% false discovery rate (FDR). Filtering was performed using an in-house linear discrimination analysis (LDA) method to create one combined filter parameter from the following peptide ion and MS^2^ spectra metrics: SEQUEST parameters XCorr and ∆Cn, peptide ion mass accuracy and charge state, in-solution charge of peptide, peptide length, and mis-cleavages. Linear discrimination scores were used to assign probabilities to each MS^2^ spectrum for being assigned correctly, and these probabilities were further used to filter the dataset with an MS^2^ spectra assignment FDR of smaller than 1% at the protein level (24). For TMT-based reporter ion quantitation, we extracted the summed signal-to-noise (S/N) ratio for each TMT channel and found the closest matching centroid to the expected mass of the TMT reporter ion. PSMs were identified, quantified, and collapsed to a 1% peptide FDR and then collapsed further to a final protein-level FDR of 1%. Moreover, protein assembly was guided by principles of parsimony to produce the smallest set of proteins necessary to account for all observed peptides. Proteins were quantified by summing reporter ion counts across all matching PSMs. PSMs with poor quality, MS^3^ spectra with more than eight TMT reporter ion channels missing, MS^3^ spectra with TMT reporter summed signal-to-noise ratio less than 100, or no MS^3^ spectra were excluded from quantification (25). The mass spectrometry proteomics data have been deposited to the ProteomeXchange Consortium via the PRIDE partner repository (26) with the dataset identifier PXD022349. Protein quantitation values were exported for further analysis in Microsoft Excel or Prism8. Each reporter ion channel was summed across all quantified proteins.

### Experimental design and statistical analysis

EVs were isolated from brain tissue of 6 female CAST.*APP/PS1* and 6 female CAST (WT) littermate control mice at 8 months of age. Statistical analysis was conducted using Prism 8 (GraphPad, Inc.). Between group comparisons were analyzed by Welch’s t-test. The Gene Ontology of identified proteins were elucidated by the Database for Annotation, Visualization and Integrated Discovery (DAVID) Bioinformatics Resources 6.8. The venn diagram and heatmap analysis were generated using Venny_2.1 (http://bioinfogp.cnb.csic.es/tools/venny/) and ClustVis (https://biit.cs.ut.ee/clustvis/).

## Results

### Biochemical and morphological characterization of EVs separated from brain tissue

We separated EVs from mouse brain tissues by ultracentrifugation and sucrose gradient ultracentrifugation as previously described (21). To check the purity of the EV preparation from mouse brain tissues, the EV fractions were analyzed for their size and number by NTA. The EV fractions were enriched the particle of small size (median 122 nm) compared to brain homogenate (median 165 nm, **Figure 1A**). The particles per protein were 2.08 × 10^7^ [particles / μg] in brain homogenate and 1.40 × 10^9^ [particles / μg] in separated EV fraction (**Figure 1B**), showing significant enrichment (p < 0.001). The EV markers such as Tumor susceptibility gene 101 protein (TSG101) and CD81 were clearly represented in EV fractions, whereas contamination markers such as 130 kDa cis-Golgi matrix protein (GM130) and cytochrome C (CYC1) in MISEV2018 guidelines (12) were absent in the EV fraction (**Figure 1C**). The separated EVs were examined by transmission electron microscopy (TEM), which shows cap-shaped morphology as commonly seen separated EVs (**Figure 1D**). These data demonstrate the successful enrichment of EV fraction from mouse brain tissues.

**Figure 1.**
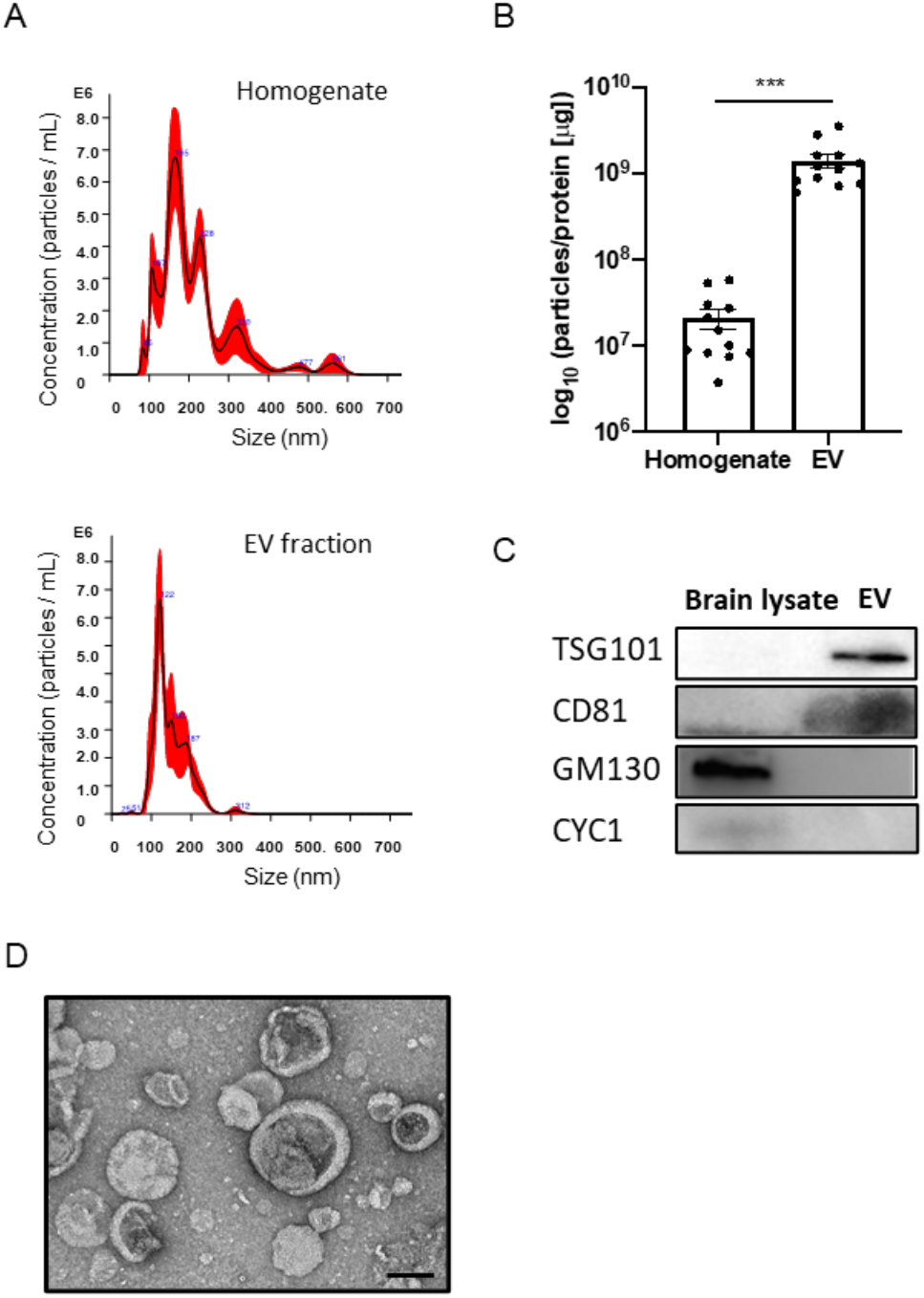
Biochemical characteristic of brain-derived EVs separated from frozen mouse brain tissue: **A)** NTA plot of average size and concentration of particles from brain homogenates and separated EV fraction. The black line shows the fitting curve. Red line represents the error bar. The y axis is the concentration of particles. The x axis is the size of particle. Top: brain homogenates, Bottom: separated EV fraction. **B)** The ratio of particles to protein concentration to quantify particle purity (*p* = 0.005 by paired samples Wilcoxon test). **C)** Assessment of EV and non-EV marker protein, including TSG101, CD81, GM130 (Golgi marker), CYC1 (Mitochondrial marker) in separated EV fraction. **D)** Transmission electron microscopy (TEM) image of mouse brain-derived EV fraction. Scale bar; 100 nm.

### Proteomic profiling of EVs from CAST.*APP/PS1* and WT mouse brain tissue

The median diameter of separated EVs was 120 nm for WT and 112 nm for *APP/PS1* groups, and the particle counts were 7.03 × 10^11^ particles for WT and 1.38 × 10^12^ particles for *APP/PS1* groups (**Figure 2A**). There were no significantly difference in these parameters between WT and *APP/PS1* groups (Diameter: *p* = 0.4848, particle counts: *p* = 0.0649). We next analyzed the protein profiles of EVs separated from *APP/PS1* and WT mouse brain tissues by LC-MS/MS/MS using TMT-based labeling (27). We identified a total of 3444 unique proteins **(Supplementary Table S1 and S2)**. The identified proteins were compared with the top 100 EV proteins from the ExoCarta database (28). The Venn diagram represents 90 of the top 100 EV proteins commonly found in the mouse brain-derived EVs **(Figure 2B)**. We analyzed the proteomic dataset using Database for Annotation, Visualization, and Integrated Discovery (DAVID Gene Ontology (GO)) (29, 30). The identified proteins show significantly enrichment of extracellular exosome by ‘cellular component’, and transport and protein-binding molecules by ‘Biological process’ and ‘Molecular function’, respectively (**Figure 2C**). The KEGG pathway analysis showed enrichment of endocytosis and glutamatergic synapse molecules, which are related to microglia and neuronal functions. The EV proteins were mostly annotated brain, brain cortex and hippocampus by Tissue ontology as expected **(Figure 2C)**. Taken together, these results show successful enrichment of proteins specific to EVs, cadherin/protein binding molecules, neuronal/glial functions and brain tissues in separated EV samples.

**Figure 2.**
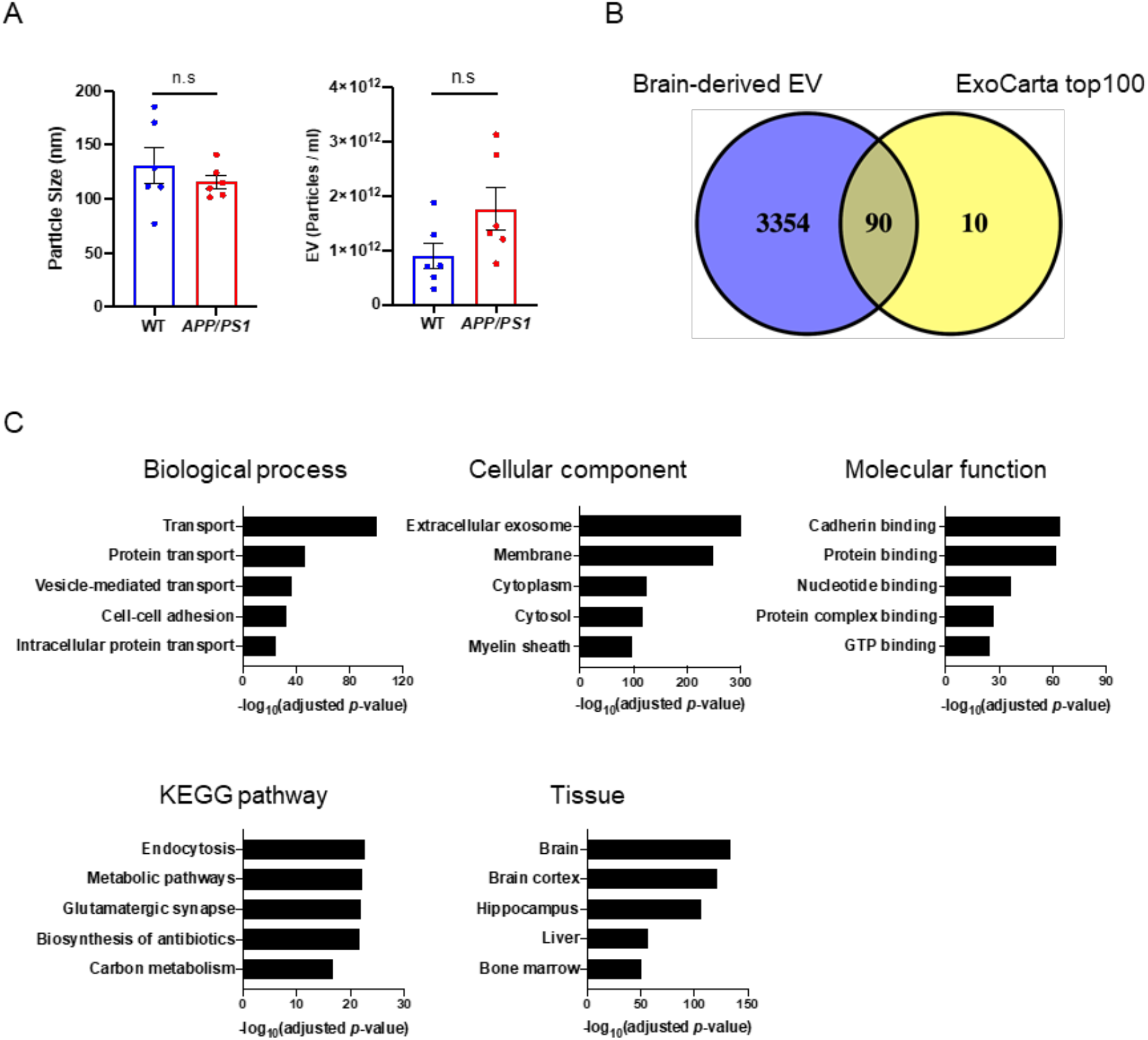
Proteomic profiling of mouse brain-derived EV: **A)** Comparison of particle number and size in EV fraction separated from CAST.*APP/PS1* and WT mouse brain tissue. Left: particle size, Right: particle number. **B)** Venn diagram representing the proteins identified in brain-derived EV and ExoCarta top100. **C)** DAVID GO analysis using DAVID Bioinformatics Resources 6.8. The GO term of Top5 Biological Process, Cellular Component, Molecular Function, KEGG pathway and Tissue ontology with −log_10_ (FDR *p*-value).

### Neural cell-type specific proteins of EVs derived from mouse brain tissue

We next examined the enrichment of neural cell-type specific molecules in the EV proteomic dataset using the proteomic dataset of neural cells, such as neurons, astrocytes, microglia and oligodendrocytes separated from mouse brain tissues by a bio-panning method as a reference (31). The identified neural cell-type specific markers (155 total) are 43.9 % (68) neurons, 5.8% (9) microglia, 27.1% (42) astrocytes and 23.2% (36) oligodendrocytes **(Figure 3A)**. We examined the changes in the expression of these cell type-specific markers in EVs separated from *APP/PS1* and WT groups. The neuron-specific molecules downregulated in *APP/PS1* compared to WT include Pclo (Piccolo), Add2 (Beta-adducin), L1cam (Neural cell adhesion molecule L1), Calb2 (Calretinin) and Calb1 (Calbindin), while upregulated molecules include Camkv (CaM kinase-like vesicle-associated protein), Gprin1 (G protein-regulated inducer of neurite outgrowth 1), Ngef (Ephexin-1), and Fxyd6 (FXYD domain-containing ion transport regulator 6) (**Figure 3B**). The astrocyte-specific molecules downregulated in *APP/PS1* compared to WT include Aldh1a2 (Retinal dehydrogenase 2), Nid1 (Nidogen-1), Lamb2 (Laminin subunit beta-2) and Cbs (Cystathionine beta-synthase), while upregulated molecules include Sorbs1 (Sorbin and SH3 domain-containing protein 1), Fmn2 (Formin-2) and Pacsin3 (Protein kinase C and casein kinase II substrate protein 3). The oligodendrocyte-specific molecules downregulated in *APP/PS1* compared to WT include P4ha1 (Prolyl 4-hydroxylase subunit alpha-1), Mog (Myelin-oligodendrocyte glycoprotein), Tnr (Tenascin-R) and Hmgcs1 (Hydroxymethylglutaryl-CoA synthase, cytoplasmic), while upregulated molecules include Col1a1 (Collagen alpha-1(I) chain), Pde9a (High affinity cGMP-specific 3’,5’-cyclic phosphodiesterase 9A) and Cnp (C-type natriuretic peptide). There are limited changes in the microglia-specific molecules identified by the previous proteomic study. To compensate the information, we have used the microglia-specific gene signature identified from microglia separated from another *APP/PS1* mouse models (32–34), namely disease-associated/neurodegenerative microglia (DAM/MGnD) and homeostatic microglia (HO). We identified DAM/MGnD-specific molecules, especially integrin alpha-x (Itgax) and apolipoprotein E (Apoe) upregulated in EVs from *APP/PS1* compared to WT group as determined by the scattered plot analysis of Log_2_ fold changes of EV proteomic dataset and the microglia gene expression profile (**Figure 3C**). These data indicate global changes in the contribution of EV production in different neural cell types, suggesting their potential application in monitoring the disease progression and understanding the pathobiology.

**Figure 3.**
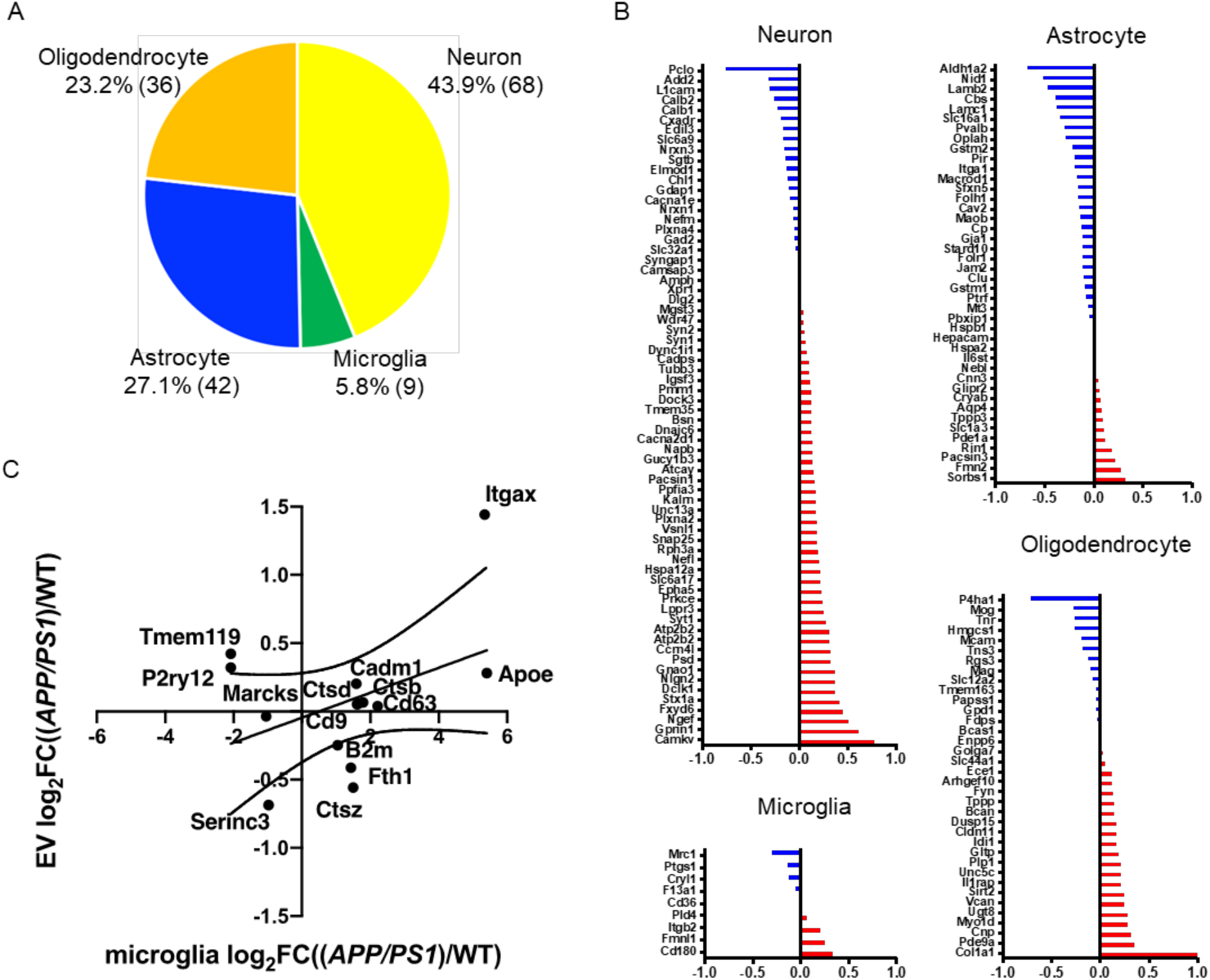
Cell type-specific proteins comparison of CAST.*APP/PS1* and WT mouse brain-derived EV: **A)** Enrichment of brain cell type-specific markers in brain-derived EV proteins. Yellow: Neuron, Green: Microglia, Blue: Astrocytes, Orange: Oligodendrocytes. The parentheses show the number of identified cell type-specific proteins. **B)** Comparison of the cell type-specific protein in CAST.*APP/PS1*-derived EVs and CAST WT EVs. The red bar shows higher expression in *APP/PS1* compared with WT and Blue bar indicates higher expression in WT compared with *APP/PS1*. **C)** Comparison of log_2_ fold change of the differential expression of DAM versus homeostatic microglia in the 5xFAD (x axis) to the log_2_ fold change of the differential expression of CAST.*APP/PS1* versus WT (y axis).

### Comparison of *APP/PS1* and WT mouse brain-derived EV proteins by TMT-labeling proteomics analysis

We analyzed the fold change and *p*-values of proteins by Volcano plot, which shows that 3 proteins were significantly upregulated (*p* < 0.05, Log_2_FC > 0.585 or < −0.5585), while 7 proteins were significantly down-regulated in *APP/PS1* compared to the WT **(Figure 4A)**. The three significantly upregulated molecules are Psen1, App and Itgax (**Figure 4B**). Among them, Psen1 and App are likely due to their transgene expression in *APP/PS1* mice, thus Itgax (CD11c), which is the most representative marker of DAM/MGnD, is the only endogenous molecule significantly upregulated in the separated EVs from *APP/PS1* mouse brain. The seven significantly downregulated molecules are WD repeat-containing protein 61 (Wdr61), Mitochondrial-processing peptidase subunit alpha (Pmpca), Retinal dehydrogenase 2 (Aldh1a2, also astrocyte-specific marker), Calumenin (Calu), Acidic leucine-rich nuclear phosphoprotein 32 family member B (Anp32b), Alpha-actinin-4 (Actn4), and NADH dehydrogenase flavoprotein 2 (Ndufv2) **(Figure 4C)**. The 46 significantly differentially expressed proteins (DEPs, *p* < 0.05) are displayed in a heatmap, showing two clusters either upregulated or downregulated in *APP/PS1* compared to WT group **(Figure 4D)**. The upregulated proteins include Anxa5 (Annexin-5), which specifically binds to the phosphatidylserine expressed on dying cells (35). We recently reported ANXA5 as the most upregulated molecules in AD brain-derived EVs compared to healthy control group (20). We also confirmed the expression of ANXA5 by immunoblotting of EVs separated from *APP/PS1* and WT mouse brains **(Supplementary Figure. S1)**. We compared the ratio of mRNA levels in *APP/PS1* mouse brain tissues over WT controls, which was published (11) and the ratio protein levels EVs separated from *APP/PS1* mouse brains over WT controls in this study by a scattered plot (**Figure 4E**). The Itgax protein show highly positive correlation with Itgax mRNA level (log_2_ mRNA expression ratio; 3.77, log_2_ EV protein expression ratio; 1.44). These data demonstrate that DAM/MGnD induction in *APP/PS1* mouse brain, as determined by Itgax expression, may contribute to the enhanced EV production by microglia, which is shown in the upregulation of Itgax in *APP/PS1* mouse brain-derived EVs. The ItgaX protein was upregulated in EVs separated from *APP/PS1* mouse brains using immunoblotting **(Figure 4F)**.

**Figure 4.**
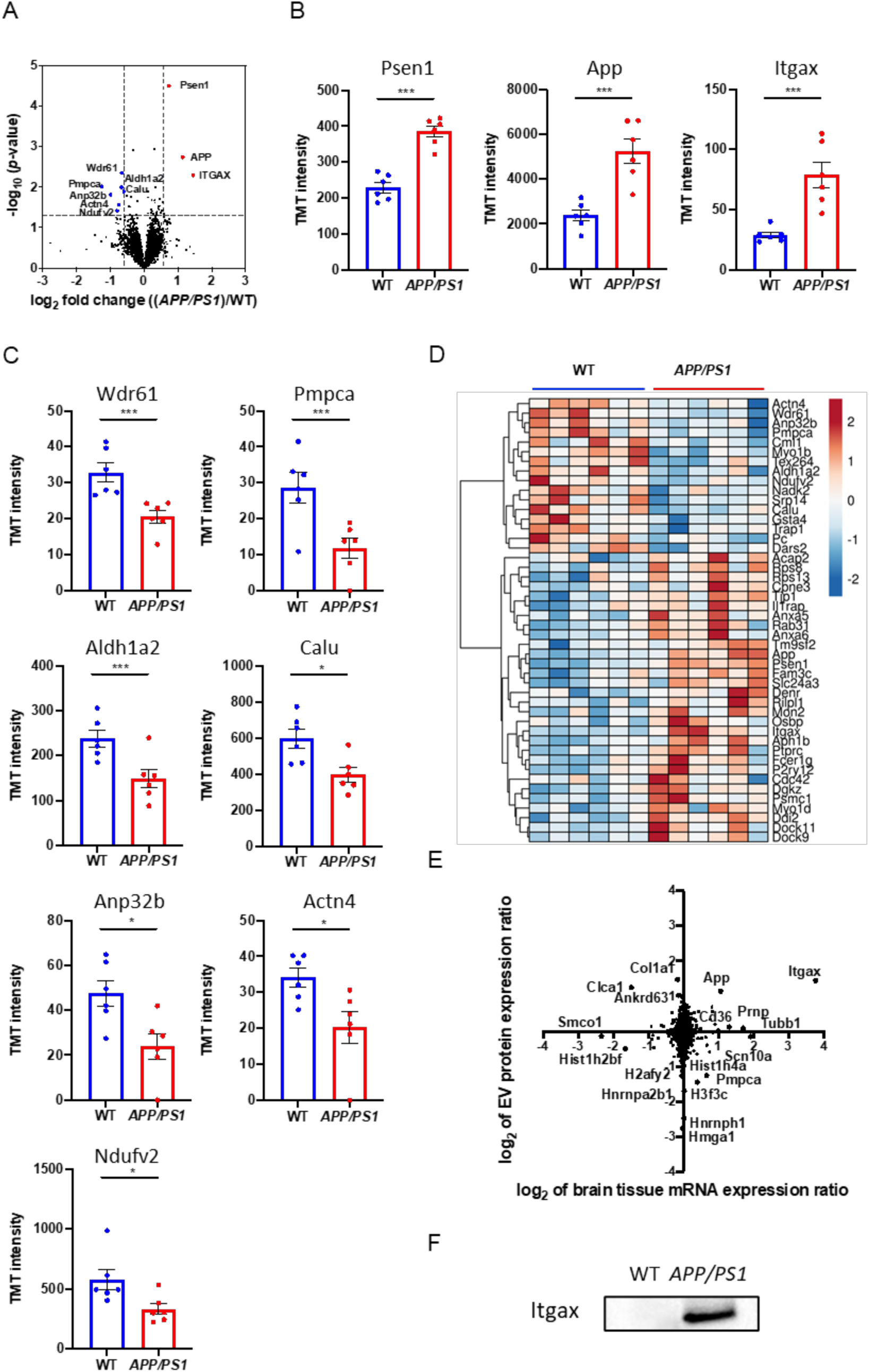
Comparison of CAST.*APP/PS1* brain-derived EV and CAST WT EV: **A)** Volcano plot showing degree of differential expression of brain-derived EV proteins in AAP/PS1 compared with WT. The x-axis indicates log_2_ transformed fold change in expression. The y-axis shows −log_10_ transformed *p*-values. The grey dot line shows the 1.3010 −log_10_(*p*-value) cutoff and 0.585 or −0.585 log_2_FC cutoff. **B, C)** A scatter plot of TMT reporter ion intensity as measured by proteomics per selected candidate protein. The t.test was calculated by Welch’s test. **B)** The three proteins were up-regulated in *APP/PS1* compared to WT. Psen1: −log_10_(*p*-value) = 4.245, FC = 1.67, App: −log_10_(*p*-value) = 2.850, FC = 2.22, Itgax: −log_10_(*p*-value) = 2.291, FC = 2.72. **C)** The 7 proteins were down-regulated in *APP/PS1* compared to WT. Wdr61: −log_10_(*p*-value) = 2.349, FC = 0.63, Pmpca: −log_10_(*p*-value) = 2.019, FC = 0.42, Aldh1a2: −log_10_(*p*-value) = 1.996, FC = 0.63, Calu: −log_10_(*p*-value) = 1.892, FC = 0.66, Anp32b: −log_10_(*p*-value) = 1.812, FC = 0.50, Actn4: −log_10_(*p*-value) = 1.562, FC = 0.59 and Ndufv2: −log_10_(*p*-value) = 1.413, FC = 0.58. **D)** Heatmap of 46 proteins with the 1.3010 −log_10_(*p*-value) cutoff. The value shows log_2_(FC). **E)** Comparison of protein expression and mRNA expression in *APP/PS1* and WT. The y axis is the ratio of EV protein expression. The x axis is the ratio of brain tissue mRNA expression. The Spearman rank correlation coefficient (rho) shows 0.06709 (*p* = 0.0001). F) Validation of Itgax in separated EV fraction from CAST.*APP/PS1* and WT mouse brain tissue using Western blot.

**Table 1.**
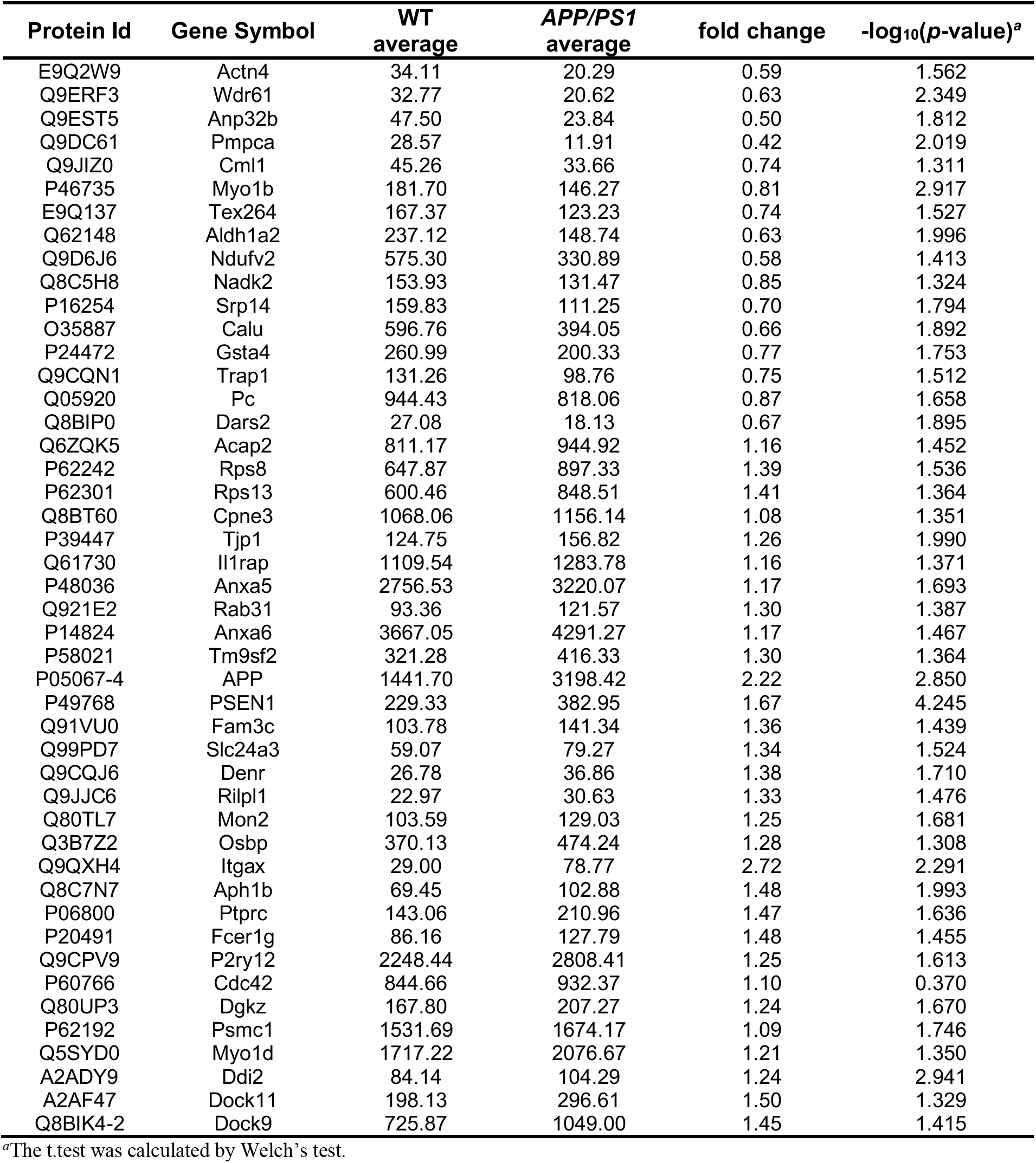

## Discussion

In the present study, we separated EVs from brain tissue of CAST.*APP/PS1* transgenic mice and age-matched CAST WT littermates. The EV samples were biophysically and morphologically characterized and subjected TMT-labeled high-resolution quantitative proteomic profiling by Nano LC-MS/MS/MS. A total of 3,444 unique proteins from brain-derived EVs, were found to be enriched as extracellular exosomes molecules. The identified EV proteins were enriched in neural cell type and DAM/MGnD -specific molecules in CAST.*APP/PS1* compared to WT group. Itgax, the DAM/MGnD marker, was significantly upregulated in EVs from CAST.*APP/PS1* compared to WT mouse brains. In addition, the significantly increased level of ANXA5 in the CAST.*APP/PS1* group, which was also increased in AD brain-derived EVs, was confirmed by western blot.

The protein levels of APP and Psen1 were also significantly upregulated in CAST.*APP/PS1* brain-derived EVs compare to WT. The peptides of APP identified in both groups by Nano-LC-MS/MS, which covered 14.5% of APP, including the corresponding amyloid-β peptide region **(Supplementary Figure S2)**. The quantification value of identified peptides, which contain amyloid beta peptide and C-terminal peptide showed to upregulate 1.5 - 2.0 folds in CAST.*APP/PS1* compared to WT in **Supplementary Table S3**. The EV in CAST.*APP/PS1*, therefore, may contain amyloid-β peptide, full-length APP and cleaved C-terminal APP. Our attempt to detect these molecules by ELISA was unsuccessful due to the scarcity of the target molecules (data not shown).

Itgax is a well-established integrin and form complex with Integrin beta2 (Itgb2/CD18) as inactivated-C3b receptor 4 (complement receptor 4) (36). The expression levels of Itgax is specifically increased in DAM/MGnD microglia separated from aged *APP/PS1* mice (37). In addition, we have recently shown that amyloid plaque-associated Mac2^+^ DAM/MGnD microglia hyper-secrete EVs to extracellular regions in *App*^*NL-G-F*^ knockin mouse model, demonstrating that DAM/MGnD plays a key role in EV secretion in AD mouse brains (38).

We compare the EV proteomics data to human AD brain-derived EV proteomics data (20), the 380 proteins were common between CAST.*APP/PS1* brain-derived EVs and human AD brain-derived EVs **(Supplementary Figure S3)**. The APOE, CAMKV, ANXA5 and VGF showed a similar correlation with these EVs, suggesting that Aβ deposition may be the major pathology for the upregulation of these molecules in EVs.

The study has some limitation. The first is the limited amount of EVs that can be separated from mouse brain (9.7 - 27.5 μg/whole brain). It is often difficult to detect proteins of interest unless highly sensitive quantification method (such as digital ELISA) is available with the limited amount of proteins. The second is the depth of identified neural cell type-specific molecules from mouse brains that are publicly available. This is especially an issue for microglia-specific markers in this study. This can be compensated by the dataset of cell type-specific gene expression analysis, but these molecules still need to be validated by proteomic approaches. Another issue is the lack of other neuropathology in *APP/PS1* mouse models, such as tau accumulation, cortical atrophy, which may attribute to the difference in proteomic profiles of EVs separated from human and mouse brain tissues. Further studies will be necessary to address these limitations by the use of more robust and sensitive protein detection systems, development of more comprehensive dataset for neural cell type-specific proteome, and application of animal models more closely recapitulating AD progression in brain.

In summary, we have profiled a total of 3,444 proteins in EV samples separated from CAST.*APP/PS1* and CAST WT mouse brain tissues at 8 months of age. *APP/PS1* mouse brain-derived EVs are enriched in App, Psen1, Itgax and Anxa5, representing the amyloid pathology progression, contribution of DAM/MGnD-derived EVs and apoptotic cell-detecting molecules: A highly relevant molecular set for understanding the disease progression in *APP/PS1* mouse brains.

## Abbreviations

Aβ: Amyloid beta peptide
AD: Alzheimer’s disease
ANXA5: Annexin-5
APOE: Apolipoprotein E
APP: Amyloid precursor protein
BCA: Bicinchoninic acid
CNS: Central nervous system
CYC1: Cytochrome C
DAM/MGnD: disease-associated / neurodegenerative microglia
DAVID: Database for annotation, visualization, and integrated discovery
DEP: Differentially expressed proteins
EVs: Extracellular vesicles
FAD: Early-onset / familial AD
GM130: 130 kDa cis-Golgi matrix protein
HO: Homeostatic microglia
ITGAX: Integrin alpha-x
LOAD: Sporadic / late-onset AD
MS: Mass spectrometry
MVBs: Multivesicular bodies
NFT: Neurofibrillary tangles
NTA: Nanoparticle tracking analysis
PS1: Presenilin-1
TEM: Transmission electron microscopy
TSG101: Tumor susceptibility gene 101 protein
UC: Ultracentrifugation
TMT: Tandem mass tag

## Acknowledgment

The authors thank M. Ericsson (Electron Microscopy Facility, Harvard Medical School) for electron microscopic imaging services.

## Author Contributions

S.M. and T.I. designed research; S.M., M.P.J., N.I., M.A., and K.O. performed research; S.M., M.P.J., N.I., J.H., and S.P.G. analyzed data; L.O., and G.H. provided brain sample; and S.M., and T.I. wrote the paper; S.M., W.P.J., N.I., M.A., S.I., and T.I. edited the paper.

