## Supplementary Figure S1-S3 for "Enrichment of neurodegenerative microglia signature in brain-derived extracellular vesicles isolated from Alzheimer’s disease mouse model"

**Muraoka S, et al.**

**Molecular & Cellular Proteomics 2020.**

**Supplementary Figures**

**Supplementary Figure S1. Assessment of ANXA5 protein by Proteomics and Western blot**

**Supplementary Figure S2. Sequence coverage of identified tryptic fragment peptide from APP by LC-MS/MS analysis**

**Supplementary Figure S3. Comparison of CAST.*APP/PS1* mouse brain-derived EV proteome and human AD brain-derived EV proteome**

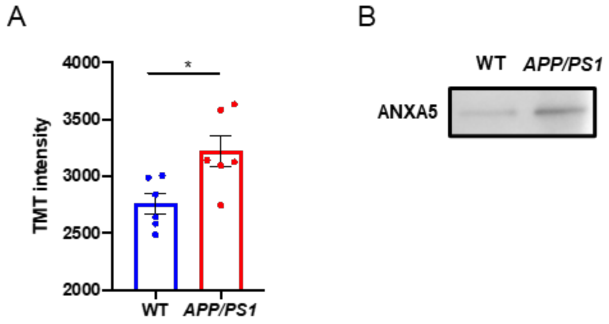


**Supplementary Figure S1. Assessment of ANXA5 protein by Proteomics and Western blot:**
**A)** A box plot of TMT-reporter intensity by proteomics. (-$\log_{10}$($p$-value) = 1.756, FC = 1.16). The t.test was calculated by Welch's test. **B)** Validation of proteomic result using Western blot.

Human Amyloid beta A4 protein (APP)

MLPGLALLLLAAWTARALEVPTDGNAGLLAEPQIAMFCGR**LNMHMNVQNGK**WDSDPSGTK
TCIDTKEGILQYCQEVYPELQITNVVEANQPVTIQNWCKRGRKQCKTHPHFVIPYRCLVG
EFVSDALLVPDKCKFLHQERMDVCETHLHWHTVAKETCSEKSTNLHDYGMLLPCGIDKFR
GVEFVCCPLAEESDNVDSADAEEDDSDVWWGGADTDYADGSEDKVVEVAEEEEVAEVEEE
EADDDEDDEDGDEVEEEAEEPYEEATERTTSIATTTTTTTESVEEVVREVCSEQAETGPC
RAMISRWYFDVTEGKCAPFFYGGCGGNRNNFDTEEYCMAVCGSAMSQSLLKTTQEPLARD
PVKLPTTAASTPDAVDKYLETPGDENEHAHFQKAKERLEAKHRERMSQVMREWEEAERQA
KNLPKADKK**AVIQHFQEKVESLEQEAANER**QQLVETHMAR**VEAMLNDR**RRLALENYITAL
QAVPPRPRHVFNMLKKYVRAEQKDRQHTLKHFEHVRMVDPKKAAQIRSQVMTHLRVIYER
MNQSLSLLYNVPAVAEEIQDEVDELLQK**EQNYSDDVLANMISEPRISYGNDALMPSLTET**
**K**TTVELLPVNGEFSLDDLQPWHSFGADSVPANTENEVEPVDARPAADRGLTTRPGSGLTN
IK**TEEISEVK**MDAEFGHDSGFEVRHQK**LVFFAEDVGSNK**GAIIGLMVGGVVIATVIVITL
VMLKKKQYTSIHHGVVEVDAAVTPEERHLSK**MQQNGYENPTYKFFEQMQN**

**Supplementary Figure S2. Sequence coverage of identified tryptic fragment peptide from APP by LC-MS/MS analysis:** Identified peptides show in Red bold. The black line indicates amyloid beta peptide.

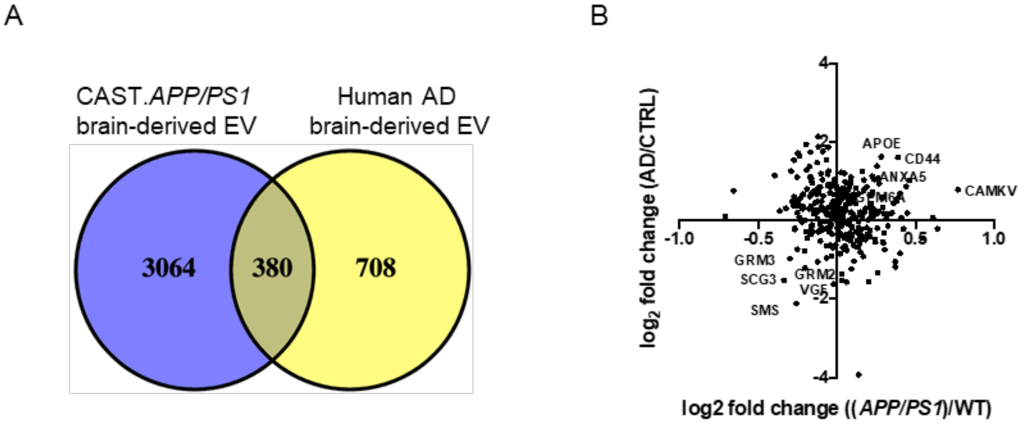


**Supplementary Figure S3. Comparison of CAST.*APP/PS1* mouse brain-derived EV proteome and human AD brain-derived EV proteome: A)** The human AD brain-derived EV proteome were identified 1088 proteins (1). The 380 proteins were common between CAST.*APP/PS1* mouse brain-derived EV and human AD brain-derived EV proteins. **B)** Scattered plot of CAST.*APP/PS1* mouse brain-derived EV and human AD brain-derived EV proteins (rho= -0.1545, $p$ = 0.0061 using two-tailed t-test).
